# Identification of differentially methylated cell-types in Epigenome-Wide Association Studies

**DOI:** 10.1101/421966

**Authors:** Shijie C Zheng, Charles E. Breeze, Stephan Beck, Andrew E. Teschendorff

**Affiliations:** CAS Key Laboratory of Computational Biology, CAS-MPG Partner Institute for Computational Biology, Shanghai Institute of Nutrition and Health, Shanghai Institutes for Biological Sciences University of Chinese Academy of Sciences, Chinese Academy of Sciences, 320 Yue Yang Road, Shanghai 200031, China; Altius Institute for Biomedical Sciences, 2211 Elliott Ave #410, Seattle, WA 98121, USA; UCL Cancer Institute, Paul O’Gorman Building, University College London, 72 Huntley Street, London WC1E 6BT, United Kingdom; Department of Women’s Cancer, University College London, 74 Huntley Street, London WC1E 6AU, United Kingdom

## Abstract

An outstanding challenge of Epigenome-Wide Association Studies (EWAS) performed in complex tissues is the identification of the specific cell-type(s) responsible for the observed differential DNA methylation. Here, we present a novel statistical algorithm, called CellDMC, which is able to identify not only differentially methylated positions, but also the specific cell-type(s) driving the differential methylation. We provide extensive validation of CellDMC on in-silico mixtures of DNA methylation data generated with different technologies, as well as on real mixtures from epigenome-wide-association and cancer epigenome studies. We demonstrate how CellDMC can achieve over 90% sensitivity and specificity in scenarios where current state-of-the-art methods fail to identify differential methylation. By applying CellDMC to a smoking EWAS performed in buccal swabs, we identify differentially methylated positions occurring in the epithelial compartment, which we validate in smoking-related lung cancer. CellDMC may help towards the identification of causal DNA methylation alterations in disease.

---

Somatic DNA methylation (DNAm) alterations have been shown to reflect cumulative exposure to environmental disease risk factors ^1^, and may also contribute to disease risk by modifying cellular phenotypes ^2,3^. One major source of DNAm variation which may hamper the identification of DNAm alterations predisposing or driving disease in Epigenome-Wide Association Studies (EWAS) ^4^, is cell-type heterogeneity ^5,6^. While statistical methods for identifying differentially methylated cytosines (DMCs) in heterogeneous tissues have been developed ^7-14^, none allow the identification of the specific cell-types responsible for the observed differential methylation ^10^. Indeed, the only existing tool that can help pinpoint differentially methylated cell-types is an enrichment analysis method for cell-type specific DNase hypersensitive sites that is performed on a relatively large list of DMCs ^15^, not allowing for individual CpGs to be ranked according to their likelihood of differential methylation (DM) in individual cell-types. Here, we present and validate CellDMC, a novel statistical algorithm that can identify interactions between phenotype and the proportions of underlying cell-types in the tissue, thus allowing for the detection of differentially methylated cytosines in individual cell-types (DMCTs).

## Results

### Detection of DMCTs with CellDMC: rationale and statistical framework

We reasoned that identification of DMCTs is possible within the same linear regression framework normally used to identify DMCs, by further inclusion of statistical interaction terms between phenotype and estimated cell-type fractions (**Fig.1a, SI fig.S1**): intuitively, if a DMC is specific to one of the cell-types in the mixture, the observed differential methylation (DM) should be most prominent when the DM analysis is restricted to samples that contain the highest fraction of that cell-type (**Fig.1b**). Therefore, in principle, these cell-type specific DM patterns can be captured through inclusion of interaction terms in the linear models (**SI fig.S1**). CellDMC analyses the DNAm patterns of interactions of all cell-types in the mixture to infer DMCTs and their directionality of change (i.e. hyper or hypomethylation) (**Fig.1**, **Online Methods, SI fig.S1**). Importantly, CellDMC works irrespectively of the number of cell-types driving DM at a given locus. For instance, in the scenario where all cell-types are uni-directionally differentially methylated to a similar degree, the level of DM should be largely independent of the fraction of any given cell-type in the sample (**Fig.1c**): statistically, there is no significant interaction between phenotype and cell-type, yet CellDMC is still able to capture these associations via classical non-interaction terms (**SI fig.S1**). CellDMC can also handle more complex scenarios, where a DMC occurs in two cell-types with opposite directionality (i.e. hypomethylated in one and hypermethylated in another) (**Fig.1d**), and which may not be identifiable by current state-of-the-art DMC calling algorithms (see later). To estimate cell-type fractions, CellDMC applies our previously validated EpiDISH algorithm ^16^ in an iterative hierarchical procedure, called HEpiDISH ^17^, which leads to improved cell-type fraction estimates in complex tissues by recognizing that cell-types are naturally arranged along a developmental tree (**Online Methods, SI fig.S2**). In the context of epithelial tissues, HEpiDISH accomplishes this by using two distinct DNAm reference matrices, a primary reference matrix for the estimation of total epithelial, total fibroblast and total immune-cell (IC) fractions, and a separate, secondary, non-overlapping DNAm reference for the estimation of underlying IC cell subtype fractions. Justification and proof that this procedure works is given elsewhere ^17^, while additional proof that HEpiDISH works is provided here in the context of identifying DMCTs. In total, HEpiDISH can estimate cell-type fractions for 9-10 different cell-types commonly found in epithelial tissues, including epithelial cells, fibroblasts and 7 major types of immune-cells (**Online Methods**) ^17^. For tissues like breast, which contain a relatively large fraction of adipocytes ^18^, the primary reference also includes a representative DNAm profile for fat cells ^17^.

**Figure-1:**
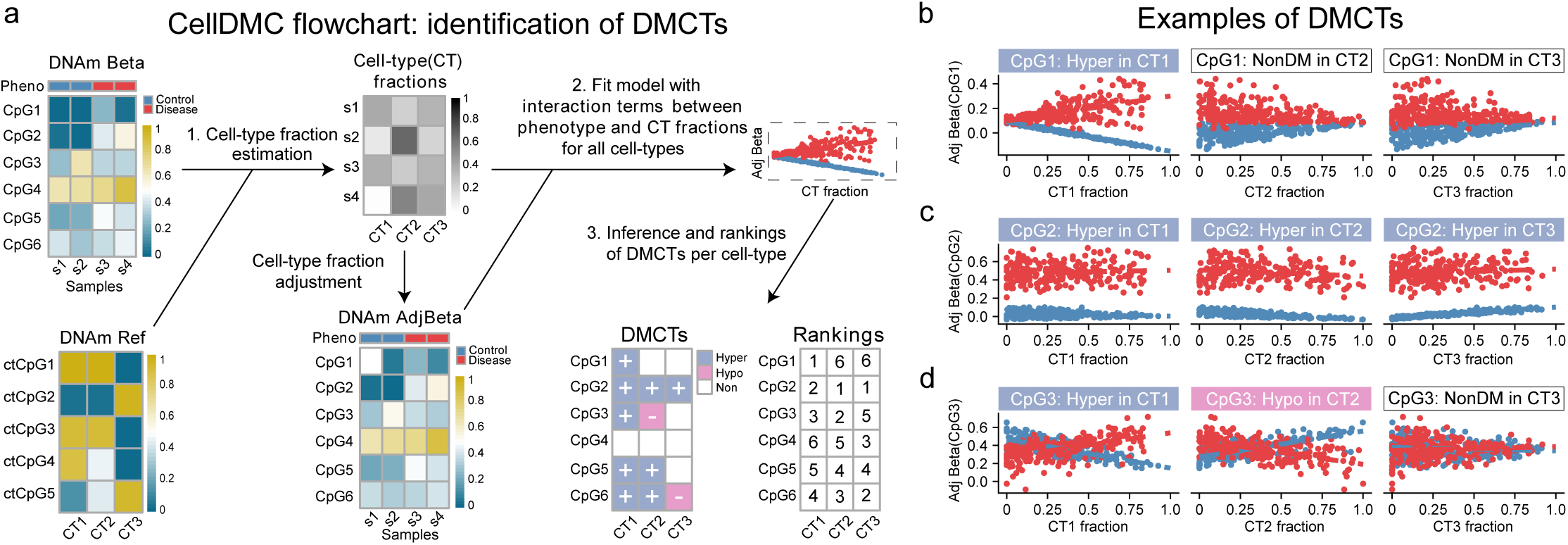
Identification of differentially methylated cell-types (DMCTs) using CellDMC. **a)** For a given DNAm data matrix, CellDMC uses a reference DNAm matrix encompassing major cell-types (CTs) in the tissue of interest, to estimate cell-type fractions in each sample, subsequently adjusting the DNAm data matrix for these estimated fractions. Using the adjusted beta-values, it then fits statistical models that include interaction terms between the phenotype and estimated cell-type fractions to identify DMCs in specific cell-types (DMCTs). These can then be ranked according to statistical significance in each cell-type. **b,c,d)** Scatterplots of adjusted beta-values against cell-type fraction for 3 different types of DMCTs. **b)** A DMCT (CpG1) which is hypermethylated in cell-type CT1 but not in cell-types CT2 and CT3. **c)** A DMCT (CpG2) hypermethylated in all three cell-types. **d)** A DMCT (CpG3) occurring in two cell-types (CT1 & CT2) but with DNAm changes occurring in opposite direction (hypermethylated in CT1, hypomethylated in CT2). Given a DNAm data matrix for an epithelial tissue that includes epithelial, stromal (fibroblast, fat) and all major immune cell types, CellDMC is able to infer not only DMCs but also the corresponding DMCTs.

### Validation of CellDMC on *in*-*silico* mixtures

To test CellDMC’s ability to detect DMCTs, we first considered a number of different simulation scenarios, where we mixed together real DNAm profiles representing epithelial, fibroblast and immune cell types, in known mixing proportions, introducing DMCs in one, two or all cell-types, with parallel or opposite directionality, and over a range of different signal-to-noise ratios (SNRs) (**Online Methods, SI table S1**). Our choice of SNRs are conservative, corresponding to differences in average DNAm within an affected cell-type of approximately 0.4 to 0.5 (i.e. 40 to 50%) in the high SNR regime (SNR~3) down to changes as small as 0.1 (i.e. 10%) for the low SNR regime (SNR<1) (**SI fig.S3**). In effect, the low SNR regime corresponds to a scenario where only 10% of say single epithelial cells exhibit a common DNAm change. Considering first scenarios where the underlying cell-type fractions do not change appreciably between cases and controls, CellDMC obtained a high sensitivity and specificity to detect DMCTs (**Fig.2a**). For instance, even when SNR<1, CellDMC obtained a sensitivity above 0.75 for all scenarios where the differential methylation is unidirectional, regardless of the number of affected cell-types (**Fig.2a**). For scenarios with bi-directional changes, CellDMC could detect DMCTs with a sensitivity above 0.75 as long as the SNR was approximately 1.8 or above (**Fig.2a**). In line with these high sensitivity values, we observed excellent agreement between the predicted DNAm difference in the underlying DMCTs with the true DNAm difference, irrespective of SNR and whether DNAm alterations were uni-or-bi-directional (**Fig.2b**). Importantly, CellDMC allows DMCTs to be ranked in each cell-type according to their statistical significance with the ranking highly correlated with true effect size, as required (**SI fig.S4**). All of the above results were largely unchanged if cell-type fractions were allowed to vary between cases and controls (**SI fig.S5**), or if DMCs only occurred in one of the underlying immune cell subtypes (compared to the previous case when the DMCs were common to all ICs) (**SI fig.S6-S8**). CellDMC also attained high sensitivity and specificity if *in*-*silico* mixtures were generated purely from IC subtypes (without admixture by epithelial and fibroblast cells), representing the more common scenario of EWAS performed in whole/peripheral blood (**SI fig.S9**), or if the phenotype is continuously valued as opposed to binary (**SI fig.S10**). CellDMC was also robust to binary heterogeneous phenotypes with cases defined by a bi-modal distribution, as observed for instance in cancer (**SI fig.S11**). We further verified that CellDMC is robust to the choice of cell-type fraction estimation algorithm (**SI fig.S12**). Underlying this robustness, we observed that results were also unchanged if cell-type fraction point estimates were perturbed using either theoretical or empirical confidence intervals (**SI fig.S13-14**).

**Figure-2:**
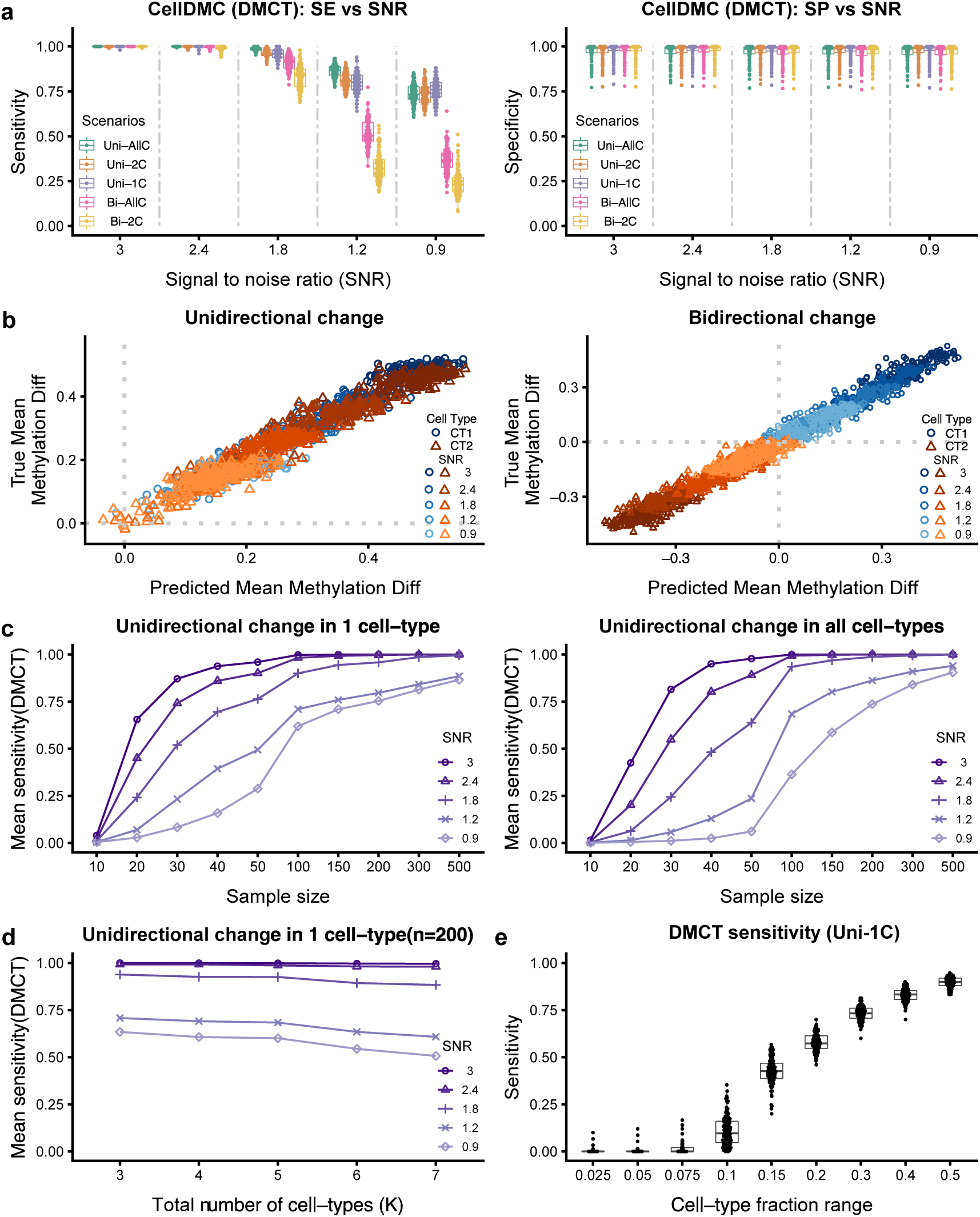
Validation of CellDMC on simulated data. **a)** Sensitivity (SE) and specificity (SP) (y-axis) of CellDMC to detect DMCTs in five different scenarios and for five different SNRs (x-axis). The scenarios correspond to uni-directional DNAm changes being unique to one cell-type (Uni-1C), shared by two cell-types (Uni-2C) or occurring in all cell-types (Uni-AllC), and to bi-directional changes (i.e hypermethylated and hypomethylated) in two cell-types (Bi-2C) or all cell-types (Bi-AllC). SNR values range from SNR~3 (mean DNAm difference in affected cell-types of approx. 0.4 to 0.5) to SNR < 1 (mean DNAm difference in affected cell-types of approx. 0.1). **b)** Scatterplots of the true DNAm difference in affected cell-types (y-axis) versus the predicted DNAm difference from CellDMC (x-axis) for the uni-directional (left panel) and bi-directional (right panel) cases. Data points are shown for 100 DMCs, 2 affected cell-types (CT1 and CT2) and for five SNR levels. **c)** Sensitivity to detect DMCTs as a function of total sample size (numbers of cases and controls are assumed equal) for five different SNR values in the Uni-1C (left panel) and Uni-AllC (right panel) scenarios. **d)** Sensitivity to detect DMCTs as a function of the total number of cell-types (K) in the Uni-1C scenario, with fractions sampled from a uniform Dirichlet distribution. In c-d) each data point represents the mean over 100 Monte-Carlo runs with 100 “control” and 100 “disease” in-silico mixtures. **e)** As d) but now as a function of the cell-fraction range exhibited by the affected cell-type in the mixture (K=3). Results shown for 100 Monte-Carlo runs.

We also assessed CellDMC’s power and specificity to detect DMCTs as a function of sample size, cell-type complexity and missing cell-types (**SI fig.S15-S18**). At a sample size of 100 controls and 100 cases, power of CellDMC was always greater than 0.75 if the SNR was larger than 1.8 (corresponding to DNAm changes greater than 0.2 in individual cell-types) (**Fig.2c**, **SI fig.S15**). Of note, as long as the fraction of the affected cell-type is distributed uniformly across samples (thus exhibiting a fairly large range), CellDMC’s sensitivity was robust to cell-type complexity, i.e. to the number K of cell-types in the mixture, although due to limitations on data availability, we could only assess this up to a value of K=7 (**Fig.2d**, **SI fig.S16**). Allowing the variance of the altered cell-type fractions to vary, we estimated that the sensitivity of CellDMC would remain over 50% as long as the range of fractions exhibited by the affected cell-type is higher than 0.2 (**Fig.2e, SI fig.S17**). CellDMC was also robust if one major unaffected cell-type was missing from the reference DNAm matrix, while reduction in sensitivity was relatively marginal if an altered cell-type was missing from the reference (**SI fig.S18**). In the case of blood tissue we simulated a realistic scenario, where cell-type fractions were modelled as observed in real blood EWAS but with one cell-type (CD8+ T-cells) missing from our reference: sensitivities to detect DMCTs in CD4+ T-cells were reduced at most by only 15% (**SI fig.S19**). Thus, these data support the view that CellDMC can reliably detect DMCTs in a wide range of different realistic scenarios. However, we also expect CellDMC’s power to be strongly influenced by cell-type complexity, in line with the fact that DNAm references become less reliable as K increases, and also because specific cell-type proportions may exhibit small variation or fall below detection levels.

### CellDMC outperforms state-of-the-art reference-based and reference-free DMC calling methods

Next, we compared CellDMC to state-of-the-art reference-based and reference-free DMC calling methods (**Fig.3, SI fig.S20**). Although CellDMC outperformed competing methods in effectively all considered scenarios, differences were especially striking when DNAm changes were bi-directional **(Fig.3, SI fig.S20-S21)**. For instance, when DNAm changes occurred in two of the underlying cell-types with opposite directionality (denoted “Bi-2C”), conventional reference-based DMC calling that only includes the estimated cell-type fractions as covariates ^7^ would not resolve the underlying DMCs, resulting in sensitivities well below 0.25 (**Fig.3b, SI fig.S20-S21**). In contrast, CellDMC attained sensitivities that ranged from over 0.95 for high SNRs to around 0.5 for low SNRs (**Fig.3a**, **SI fig.S20-S21**). If bi-directional changes occurred in more than 2 cell-types (denoted “Bi-AllC”), CellDMC also attained significantly higher sensitivity values than conventional non-interaction based DMC calling (**Fig.3a-b, SI fig.S20-S21**). Similar improvements in sensitivity of CellDMC over state-of-the-art reference-free methods (SVA & RefFreeEWAS) ^8,19,20^ were also observed, which were again particularly prominent in bi-directional scenarios affecting two or more cell-types (**Fig.3a,c-d**). In line with the sensitivity results, CellDMC’s specificity was also higher and more stable than that of competing methods (**Fig.3e-h, SI fig.S20**). Thus, these data demonstrate that CellDMC can attain sensitivity and specificity values close to 100% in scenarios where current state-of-the-art methods would fail to detect DMCs.

**Figure-3:**
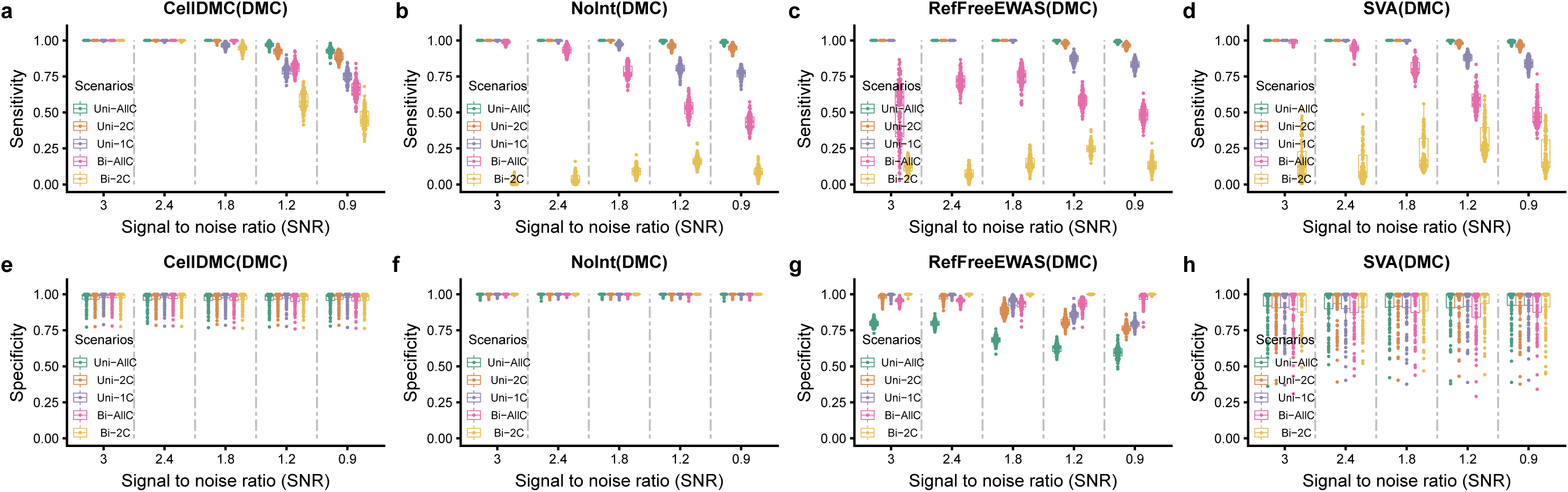
Benchmarking of CellDMC to state-of-the-art reference based and reference-free DMC calling algorithms. **a-d)** From left to right, plots of the sensitivity of CellDMC, of reference-based DMC calling (which includes cell-type fractions only as covariates without interaction-terms denoted “NoInt”), of reference-free methods RefFreeEWAS and SVA, against the SNR for five different differentially methylated cell-type scenarios, where mean cell-type fractions are not different between cases and controls. Because the competing methods can only detect DMCs, and not DMCTs, we compare the methods in terms of their sensitivity to detect DMCs. Results are for 100 Monte-Carlo runs, where in each run 200 in-silico mixture samples were simulated (100 “control” and 100 “disease”). Note that for the bi-directional scenario, competing state-of-the-art methods cannot reliably detect DMCs, because the opposite directional DNAm change in two of the underlying cell-types (assumed here to be of equal magnitude) “cancel-out”. **e-h)** As a-d), but now for the specificity.

### Validation of CellDMC on whole-genome bisulfite sequencing data

All previous simulation models mixed together DNAm profiles generated with Illumina 450k/EPIC technology ^21,22^, a platform matched to the one used to generate the DNAm references. Thus, to demonstrate applicability of CellDMC to samples generated with a different technology, we devised *in*-*silico* mixture simulation models using whole-genome bisulfite sequencing (WGBS) profiles of purified cell-types, generated as part of IHEC ^23^ (**SI fig.S22**). Although CellDMC’s performance measures dropped upon application to WGBS data, the reduction was only relatively marginal, with sensitivity and specificity values remaining high (sensitivity and specificity was still over 90% for the higher SNR values, **SI fig.S23**). Thus, we conclude that profiling technology is not a major limiting factor for CellDMC.

### Validation of CellDMC in a blood EWAS

Next, we tested the ability of CellDMC to detect known DMCTs in a real EWAS. A recent EWAS for Rheumatoid Arthritis (RA) conducted on purified B-cells identified a total of 10 DMCs, which were validated in two independent EWAS cohorts which also profiled purified B-cells ^24^ Thus, we reasoned that application of CellDMC to an independent RA EWAS performed on 689 blood samples ^5^, should be able to identify these 10 RA-DMCs as being differentially methylated specifically in B-cells. CellDMC predicted 8 out of these 10 to be differentially methylated, and 7 out of 10 to be B-cell DMCTs (FDR<0.05, **Fig.4a**). Of note, CellDMC predicted the DNAm difference between RA cases and controls to be occurring primarily in B-cells, and not in the other immune cell subtypes (**Fig.4a**), despite the fact that on average B-cells only accounted for 2% of blood cell subtypes in the samples, and despite exhibiting relatively little variation (**Fig.4a**). Although the median absolute difference in B-cell fractions between the 689 samples was only 1.5%, 5 samples exhibited fractions larger than 10%, and therefore the ability of CellDMC to detect validated B-cell DMCTs is consistent with our previous simulation estimates (**Fig.2e**). Importantly, we estimated the chance that CellDMC would call 7 out of 10 randomly selected CpGs to be B-cell DMCTs at their observed significance levels, to be less than 0.00001 (**Fig.4a**). We verified that results were robust to the choice of cell-type fraction estimation algorithm (**SI fig.S24**) or if one of the other minor blood cell subtypes was removed from the DNAm reference matrix (**SI fig.S25**). Of note, had we applied a standard non-interaction based reference model to identify DMCs, 6 of the 10 validated RA-DMCTs would have been detected as DMCs using the same FDR<0.05 threshold but without knowledge of the underlying DMCT being B-cells. We also note that CellDMC led to a larger than 4-fold reduction of DMCs compared to a method that did not adjust for cell-type fractions, and a larger than 2-fold reduction of DMCs compared to a non-interaction based model, with relatively little overlap between DMCs called by CellDMC and the standard non-interaction model (**SI table.S2**). Thus, like the standard non-interaction based model, CellDMC is able to remove large numbers of associations caused by changes in the granulocyte/lymphocyte ratio ^5^, whilst also identifying different DMCs to those found using the standard model.

**Figure-4:**
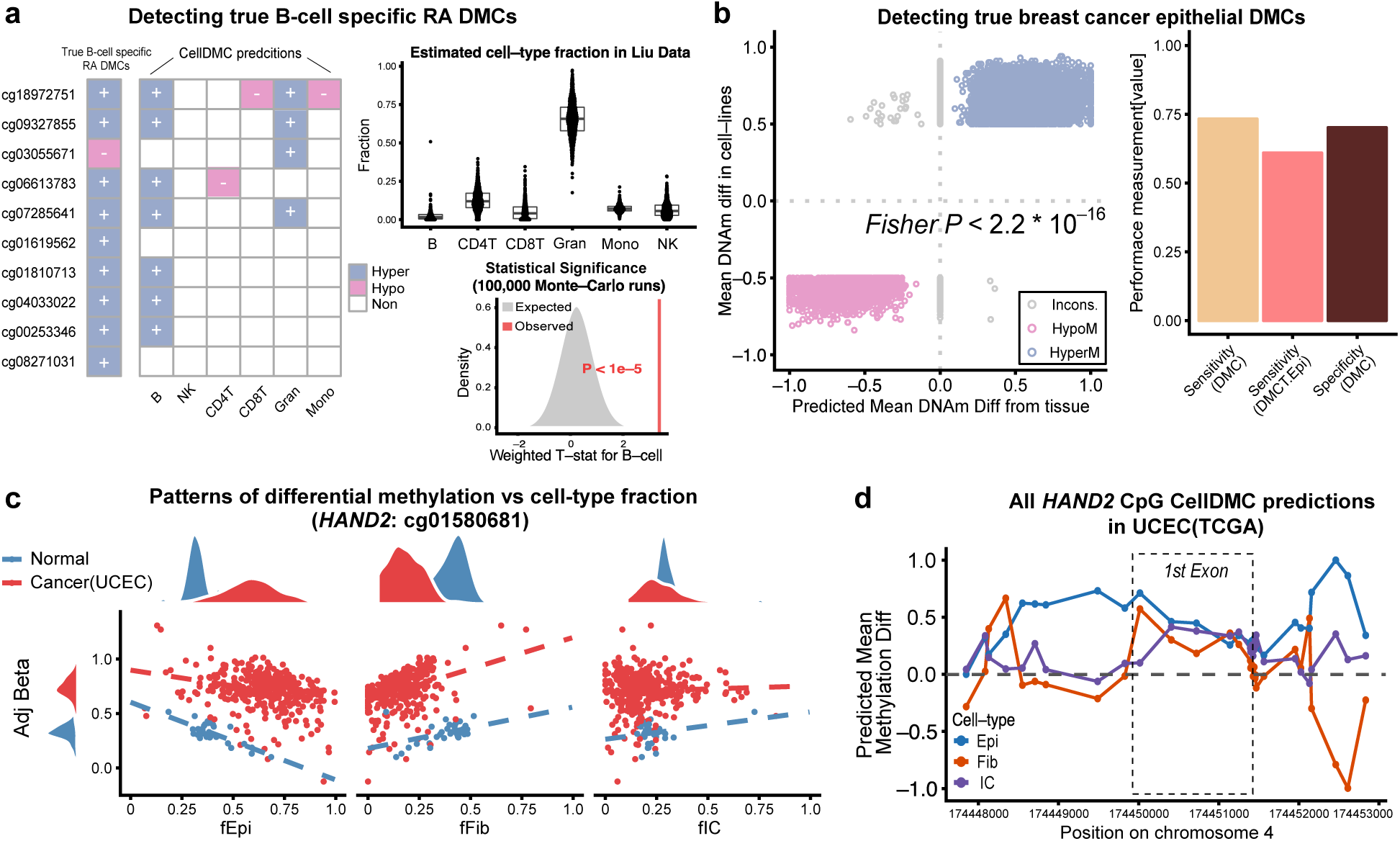
Validation of CellDMC in real EWAS and cancer epigenome data. **a)** Validation of CellDMC in a Rheumatoid Arthritis (RA) EWAS performed on 689 blood samples (Liu et al ^5^). The bar to the left indicates 10 CpG sites that have been validated to be RA-associated DMCs in 3 independent purified B-cell cohorts, with their directionality of DNAm change as indicated ^24^ The central heatmap displays the predicted DMCTs from CellDMC. Statistical significance of this result was assessed using 100,000 Monte-Carlo runs, where in each run 10 CpGs were selected at random and their weighted averaged t-statistic (grey curve and area) compared to the observed value (vertical red line). Boxplots compare cell-type fraction estimates in the 689 samples between cell-types. **b)** Validation of CellDMC to detect true breast cancer epithelial DMCs. Scatterplot of the DNAm difference between breast cancer cell lines and normal mammary epithelial cell-lines (y-axis) against the predicted DNAm difference from CellDMC for the epithelial subfraction of breast cancer tissue compared to the epithelial subfraction of normal breast, for a total of approximately 19,000 true positive breast cancer epithelial DMCs. Fisher-test P-value is given. Barplots to the right display CellDMC’s sensitivity to detect DMCs and DMCTs as well as specificity. **c)** Validation of CellDMC in the endometrial TCGA cancer study. Scatterplots of the beta value (adjusted for cell-type fractions) of one of the *HAND2* CpGs against the epithelial, fibroblast and total immune-cell (IC) fractions, with samples labeled according to endometrial cancer (red) or normal endometrium (blue). Right panel displayss the predicted DNAm difference between cancer and normal in each of the 3 cell-types, for all CpGs mapping to *HAND2.*

### Validation of CellDMC in cancer epigenome studies

To demonstrate the ability of CellDMC to detect DMCTs in solid epithelial tissues, we applied it to the breast cancer EWAS setting, for which we had previously constructed a gold-standard set of 19,379 true positive breast cancer epithelial DMCs (bcDMCs) ^9,17^ This set was obtained by intersecting DMCs from a comparison of breast cancer epithelial to normal mammary epithelial cell-lines (thus representing relatively pure epithelial cell populations), with a corresponding list of DMCs derived from the TCGA breast cancer study ^17,25^. We reasoned that applying CellDMC on our independent breast cancer tissue EWAS ^18^ should not only validate the breast cancer epithelial DMCs, but should also predict the epithelial compartment to be the main DMCT. Confirming this, CellDMC’s predicted mean methylation differences between the epithelial cells found in breast cancer and those found in the normal tissue, exhibited an excellent correlation with the DNAm differences observed in the actual cell-lines (**Fig.4b**). Overall, the magnitude of DNAm changes were still larger in the cell-lines possibly owing to cell-culture effects or their longer proliferative history (**Fig.4b**). CellDMC exhibited approximately 75% sensitivity and 70% specificity to detect DMCs and over 60% sensitivity to detect the DMCTs as occurring in the epithelial compartment (**Fig.4b, SI fig.S26a**). Although it is unknown if the 19,379 true positive bcDMCs are also altered in fibroblasts, fat and immune cells, CellDMC predicted much smaller fractions of these loci to be altered in fat and ICs (**SI fig.S26b**).

As a third validation on real data, we considered the case of the *HAND2* gene in endometrial cancer ^26,27^ We had previously discovered and validated hypermethylated DMCs in endometrial cancer, mapping to the 1^st^ exon region of the *HAND2* gene, which is a main target of the progesterone receptor tumor suppressor pathway ^26,27^ *HAND2* is expressed in normal endometrial tissue including stromal fibroblasts, but lacks expression in endometrial cancer and the stromal fibroblasts of the cancer tissue ^26^, suggesting that the observed hypermethylation occurs in both epithelial and fibroblast cells. Indeed, the relatively large difference in DNAm (Δβ~0.5) observed between endometrial cancer and normal endometrial tissue ^26^ is a strong indication that the unidirectional DNAm change is occurring in both epithelial and fibroblast compartments. Application of CellDMC to the TCGA endometrial cancer study ^28^ confirmed that the DNAm change is occurring in the epithelial and fibroblast cells of the tissue (**Fig.4c-d**), and that the alteration in fibroblasts is local to the 1^st^ exon region (**Fig.4d, SI fig.S27**). Interestingly, CellDMC also predicted the immune-cell compartment to be differentially methylated, albeit less strongly so (**Fig.4c-d**). This is also consistent with the observation that these *HAND2* CpG sites become hypermethylated with age in blood ^29^, and that the unmatched TCGA endometrial cancers were derived from older women. Thus, the prediction of CellDMC that these specific *HAND2* CpGs define DMCs in all 3 major cell-types within the endometrial tissue is entirely consistent with previous knowledge and data.

### CellDMC identifies smoking-associated DNAm changes in squamous epithelial cells

Finally, to demonstrate how CellDMC may help gain novel biological insight, we applied it to the largest EWAS cohort performed in buccal swabs ^30^, consisting of 790 samples from women all aged 53 but with a wide range of different lifetime levels of smoking exposure. As expected, buccal swabs contained mainly epithelial and immune cells (**SI fig.S28**), and thus we used CellDMC to predict smoking-associated DMCTs in these two cell-types. By comparison to a gold-standard list of 62 true smoking-associated DMCs derived from whole blood EWAS (validated in at least three independent whole blood EWAS studies) ^31^, we confirmed CellDMC’s sensitivity to detect true DMCTs: most of the gold-standard DMCs were predicted to be hypomethylated specifically in the immune cell compartment of the buccal tissue (**Fig.5a-d)**, in line with their observed hypomethylation in blood EWAS ^31^. Confirming CellDMC’s specificity, the top ranked DMCTs in immune cells have all been previously validated as being smoking-associated DMCs in blood (**Fig.5a)**. Although most of the 62 gold-standard smoking-DMCs in blood were not predicted to be altered in the epithelial cells of the buccal swabs, the great majority (i.e over 90%) of DMCTs were, however, predicted to occur in the epithelial compartment, with a strong skew towards hypermethylation (**Fig.5b)**. To validate these epithelial DMCTs, we posited that their variance in DNAm levels would increase with the fraction of epithelial cells in lung squamous cell carcinoma ^32^, a cancer strongly associated with smoking, whilst also exhibiting a concomitant decrease in variance in samples with a higher immune-cell content. We were able to confirm this pattern for the top-ranked hypomethylated epithelial DMCT (**Fig.5e**), and for most of the other epithelial DMCTs, at high statistical significance (**Fig.5f, Online Methods**). Moreover, predicted hyper-and-hypo methylated epithelial DMCTs exhibited a highly significant trend towards positive and negative correlations between their DNAm levels and the estimated epithelial cell-type fractions, as required (**Fig.5f**). To confirm the biological and clinical significance of these findings, we posited that the DNAm alterations happening in the epithelial cells would exhibit increased deviations in lung cancer compared to the normal adjacent tissue, owing to an increase in the epithelial fraction in the tumors (**Fig.5g**). Confirming our expectation, hypermethylated and hypomethylated epithelial DMCTs exhibited increased and decreased levels of DNAm in lung cancer, respectively (**Fig.5h**). Consistent with the total immune cell fraction not being altered between normal and cancer, the top ranked DMCTs specifically hypomethylated in immune cells did not exhibit decreased levels in cancer, but increased levels, probably due to shifts in the specific immune cell subtype proportions (**Fig.5h**) Thus, the epithelial smoking-DMCTs identified here represent epigenetic alterations occurring in the squamous cell of origin of smoking related lung cancer and therefore may mark cells that are being selected for during lung carcinogenesis.

**Figure-5:**
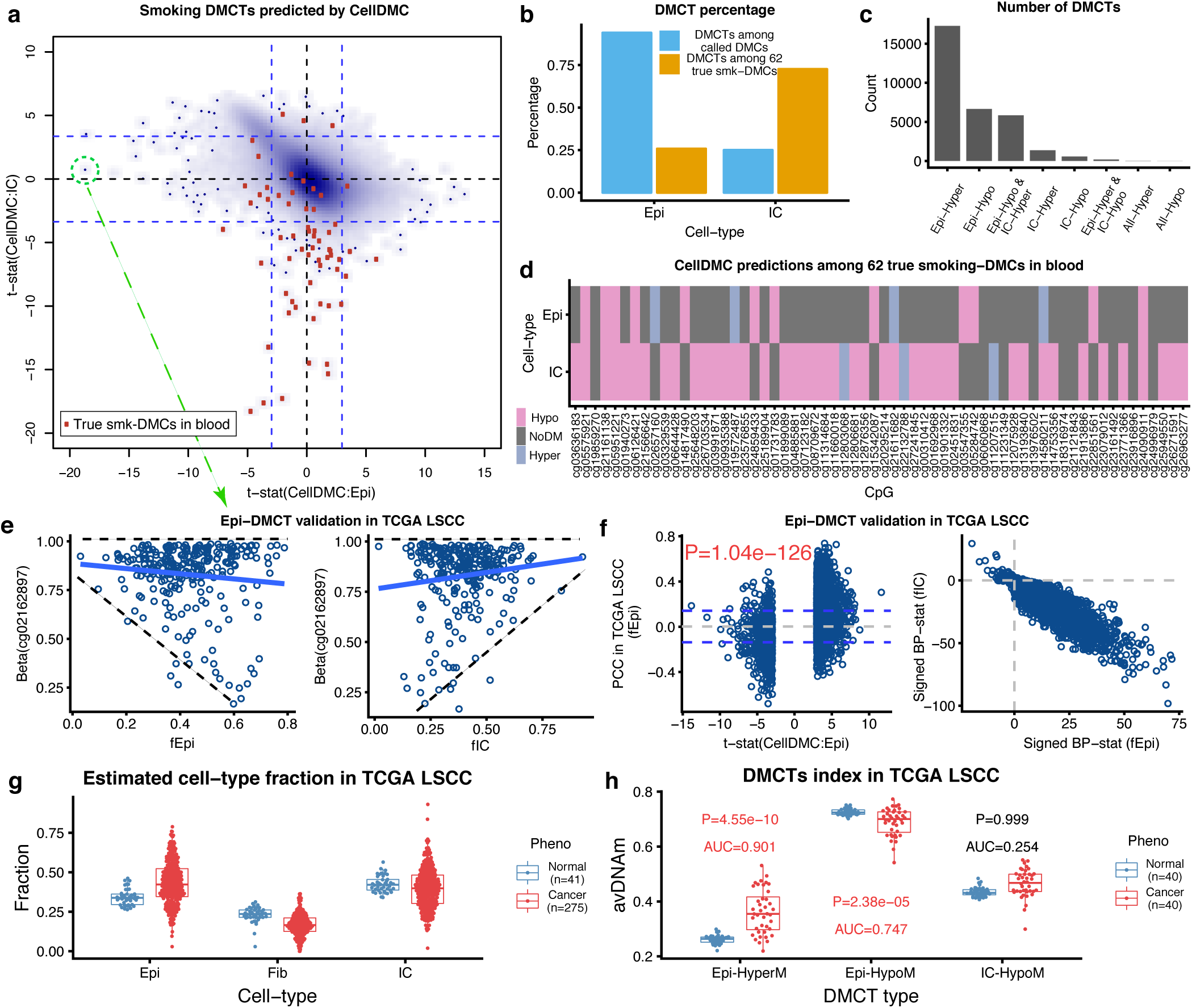
Identification of epithelial specific smoking DMCTs and their relevance in lung cancer. **a)** Smoothed scatterplot of CellDMC t-statistics in epithelial cells (x-axis) vs immune cells (IC) (y-axis) derived from a 790 buccal swab EWAS with smoking as the phenotype. Blue dashed lines indicate level of statistical significance (FDR<0.05). Red points label CpGs that belong to a gold-standard list of 62 true smoking-DMCs in blood. **b)** Barplots comparing the percentage of DMCTs that occur in epithelial and IC compartments, measured relative to the set of 62 smoking-DMCs in blood, or relative to all called DMCs/DMCTs. **c)** Barplots displaying the number of different types of DMCT, as indicated. Epi=epithelial only, IC=immune cell only, Hyper=hypermethylated in smokers, Hypo=hypomethylated in smokers. All means a DMC that occurs in both cell-types. **d)** Heatmap of CellDMC predictions (significance of signed adjusted P-values) for the 62 true smoking-DMCs. **e)** Scatterplot of the DNAm level (beta-value) of the top-ranked smoking-associated epithelial-specific DMCT (hypomethylated) against the epithelial cell-type fraction (left panel) and total immune cell fraction (right panel) across the 275 lung squamous cell carcinoma samples from the TCGA. Observe the trend towards hypomethylation and increased variance as epithelial fraction increases. **f) Left panel:** Scatterplot of the Pearson Correlation Coefficients (PCC, y-axis) between DNAm and epithelial cell-type fraction, as computed over the lung cancer TCGA samples, against the CellDMC t-statistics for predicted epithelial-specific DMCTs derived in the buccal swab smoking EWAS. Dashed lines indicate level of statistical significance and P-value is from a one-tailed Fisher’s exact test on the CpGs passing significance in each quadrant. **Right panel:** Scatterplot of the signed Breusch-Pagan (BP) differential variance statistic of DNAm against immune-cell fraction (y-axis) versus the corresponding statistic for the epithelial fraction, as computed across the TCGA lung cancer samples, for all predicted epithelial-specific smoking-DMCTs derived from buccal swab EWAS. **g)** Boxplots of cell-type fractions estimated with EpiDISH for epithelial, fibroblast and immune cells in the normal and lung squamous cell carcinoma (LSCC) samples from the TCGA. **h)** Average DNAm levels of three categories of DMCTs, derived with CellDMC from the buccal smoking EWAS, in the matched normal-adjacent and LSCC samples from the TCGA. The DMCT categories are the 50 top-ranked hypomethylated and hypermethylated (promoter) epithelial DMCTs, and the top-ranked 50 hypomethylated immune-cell DMCTs. P-values are from a one-tailed Wilcoxon rank sum test.

## Conclusions

In summary, we have presented a novel algorithm aimed at identifying DMCTs, and have extensively validated the method on simulated as well as real data, encompassing common EWAS and cancer epigenome scenarios, and different technologies. The ability of CellDMC to rank CpGs according to their probability of being DM in each of the underlying cell-types will be invaluable for EWAS and cancer epigenome studies to help improve biological interpretation, for prioritizing candidates that require functional validation, and to help elucidate causal pathways to disease.

### Software Availability

CellDMC is freely available as a user-friendly R-script within the EpiDISH Bioconductor package, freely available from github (https://github.com/sjczheng/EpiDISH), and in due course also from Bioconductor (https://www.bioconductor.org).

## Author Contributions

Study was conceived by AET. SCZ and AET contributed jointly to methods development. SCZ performed the statistical analyses. Manuscript was written by both SCZ and AET. SB and CEB contributed valuable feedback.

## Competing Interests

The authors declare that they have no competing interests.

## Acknowledgements

This work was supported by NSFC (National Science Foundation of China) grants 31571359, 31771464 and 31401120, a Wellcome Trust grant (99148) and by a Royal Society Newton Advanced Fellowship (NAF project number: 522438, NAF award number: 164914).

## Online Methods

### Estimation of cell-type fractions in complex tissues using EpiDISH/HEpiDISH

Estimation of the fractions of cell-types in a given sample constitutes the first step of the CellDMC algorithm and uses our previously validated EpiDISH ^16^ and HEpiDISH ^17^ methods. We use these methods to estimate cell-type fractions for *in*-*silico* generated mixtures as well as for real samples from EWAS and cancer epigenome studies. Which method we use, i.e. EpiDISH or a hierarchical iterative version of it called HEpiDISH depends on the desired cell-type resolution, as we explain next.

In the case of whole blood, peripheral blood or *in*-*silico* mixtures of blood cell subtypes, we use the EpiDISH procedure with a DNAm reference matrix defined over 333 IC cell subtype specific DMCs and 7 IC cell subtypes ^16^ Briefly, EpiDISH models the DNAm profile of any given sample as a linear mixture over the DNAm profiles for the individual cell-types making up the sample. The associated multivariate linear regression is run using only the 333 IC cell subtype specific DMCs, and is performed using a robust estimator, hence the specific implementation we use is Robust Partial Correlations (RPCs) ^16^ In this framework, the non-negativity and normalization constraints on the cell-type fractions (i.e. the estimated regression coefficients) are imposed *a*-*posteriori*, i.e. negative weights are set to zero and all other non-zero weights are normalized so that their sum adds to 1. As shown by us ^16^, RPCs improves inference over the alternative approach which is to impose these constraints during inference via quadratic programming optimization ^7^. RPCs also performs similarly to another non-constrained approach that uses a penalized support vector regression model called CIBERSORT ^16,33^. Hence, for these reasons, we here use EpiDISH with RPCs.

In the case of solid epithelial tissues and *in*-*silico* mixtures of epithelial, fibroblast and various IC cell subtypes, and when we only need to estimate cell-type fractions for the total epithelial, total fibroblast and total IC fractions, we once again use EpiDISH with RPCs, but now with a DNAm reference matrix defined over 716 DMCs and 3-cell types (generic Epithelial, generic Fibroblast, and a generic IC, denoted as the EpiFibIC reference matrix), as derived and validated previously by us in Zheng et al ^17^. The 716 DMCs are highly discriminative of the 3 main cell-types, exhibiting large differences in mean DNAm between epithelial, fibroblast and ICs ^17^. In some of the simulation scenarios, we also estimate fractions for individual IC cell subtypes, in which case we use our validated HEpiDISH algorithm ^17^. As explained in Zheng et al, HEpiDISH works by implementing EpiDISH in an iterative hierarchical fashion, first estimating total epithelial, total fibroblast and total IC fractions using the EpiFibIC reference, and then estimating proportions for the 7 individual IC cell subtypes using a 188 DMC subset of the original 333 DMC reference used to estimate fractions in blood. A key point is that the 188 DMCs do not share any overlap with the 716 DMCs of the EpiFibIC reference, and that all 188 DMCs have been selected to ensure that their median DNAm levels do not vary appreciably (most differences are less than 0.05 and the maximum difference is 0.3, in absolute terms) between the epithelial, fibroblast and the collection of ICs ^17^. This latter criterion helps ensure that the error or noise induced by inferring IC cell subtype fractions in the mixture using a reference matrix that does not include epithelial and fibroblast references, is less than the effect size or signal of the 188 DMCs (since any of the original 333 DMCs in the blood reference exhibit big differences in DNAm, i.e. at least +/−0.5 with the great majority exhibiting over +/−0.8 differences between at least one IC cell subtype and all others. See Zheng et al ^17^ for further justification and validation.

### The CellDMC algorithm

CellDMC consists of 3 basic steps: (1) Estimation of cell-type fractions in a given sample using EpiDISH ^16^ or Hierarchical EpiDISH (HEpiDISH) ^17^, depending on the cell-type resolution desired (as described in previous section), (2) Estimation of differentially methylated cell-types (DMCTs) for a given phenotype of interest (binary or continuous), (3) Ranking of DMCTs per cell-type.

The key idea behind CellDMC is that a DNAm alteration occurring in a specific cell-type will exhibit a significant interaction with the corresponding cell-type fraction variable. To introduce relevant notation, we shall assume that we have a DNA methylation beta-valued matrix *x_cs_*, with *c* (*c* = 1,..,*C*) labeling the CpGs and *s* (*s* = 1, …,*S*) labeling the samples. For each sample *s*, let *y_s_* denote the phenotype (e.g. a 0 or 1 variable for binary phenotypes, or a continuously-valued number for continuous phenotypes), and let
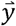
denote the vector of phenotype values over all samples. Correspondingly, we define
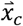
to be the vector of beta (DNAm) values for CpG *c* across the *S* samples. We further assume that the fractions for the main cell-types, indexed by *k* (*k*=*1*…*K* and with *K* the number of cell-types), in the given samples have already been estimated using EpiDISH/HEpiDISH (or another algorithm the user wishes to use). We denote the estimated fractions for cell-type *k* over the *S* samples with the vector
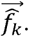
CellDMC is then based on the following statistical model, which is run separately for each CpG *c*:

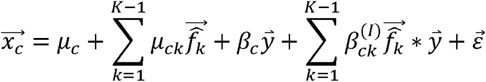

where
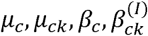
are regression coefficients to be estimated, ^∗^ denotes the interaction term and where we assume that errors are Gaussianly distributed with a variance that may depend on the specific CpG *c.* We note that the above summations don’t run over all *K* cell-types, because cell-type fractions are normalized and must add to 1. Because of this normalization constraint, the intercept term and the phenotype main effects term can be absorbed into the corresponding summations, so that an equivalent statistical model is

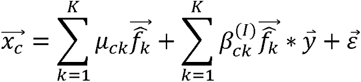

The regression coefficients
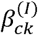
inform us as to whether there is a significant interaction between the phenotype and the corresponding fraction for cell-type *k*. We note that if differential methylation associated with the phenotype occurs at a CpG *c* and in cell-type *k* that the observed differential methylation should be larger in samples with high fractions for that cell-type *k* compared to samples with low content for cell-type *k*, and should be detectable via a statistically significant interaction term
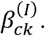
We solve the above model using least squares with the *lm* function in R, which provides estimates for the regression coefficients and their statistical significance via P-values
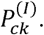. The P-values
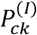
for each cell-type *k* are adjusted for multiple hypothesis testing using Benjamini-Hochberg (BH) FDR estimation. For those CpGs with BH-adjusted P-values less than a predefined significance threshold (i.e. typically BH FDR<0.05), we call it a DMCT in the given cell-type. Finally, CpGs can be ranked within each cell-type according to the associated P-value of significance.

Finally, we note that additional covariates representing other biological (e.g. age, gender, ethnicity) or technical factors (batch) can be included in the above model. For instance, for a set *Q* of such factors, let
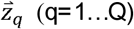
denote the factor values across the *S* samples, then the model above becomes

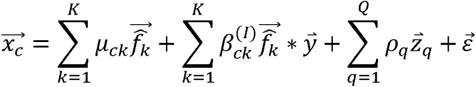

### DNA methylation datasets used for *in*-*silico* mixture simulation experiments

The following lists the DNAm datasets used in this manuscript to generate the *in*-*silico* mixtures, with their GEO (www.ncbi.nlm.nih.gov/geo) accession numbers or download links. We note that all these samples were not used in the construction of the DNAm references used in EpiDISH and HEpiDISH and therefore represent truly independent samples. Illumina 450k data of 4 epithelial cell-lines and 10 fibroblasts cell-lines from Stem-Cell Matrix Compendium-2 (SCM2) ^34^ (GSE31848), Illumina 450k data of 8 B-cells, 71 CD4+ T-cells, and 28 monocytes from Absher et al ^35^ (GSE59250), Illumina 450k data of 31 CD4+ T-cells from Limbach et al ^36^ (GSE71955), Illumina 450k data of 4 B-cells, 6 CD4+ T-cells, and 5 monocytes from Mamrut et al (GSE71244), Illumina 450k data of 36 monocytes from Marabita et al ^37^ (GSE43976), Illumina 450k data of 8 CD4+ T-cells from Nestor et al ^38^ (GSE50222), Illumina 450k data of 214CD4+ T-cells, and 1202 monocytes from Reynolds et al ^39^ (GSE56047), Illumina 450k data of 6 B-cells, 6 CD4+ T-cells, and 6 monocytes from Zilbauer et al ^40^ (https://www.ebi.ac.uk/arrayexpress/experiments/E-MTAB-2145/). Two fibroblasts cell-lines were removed from SCM2 dataset after purity check. All of the above Illumina 450k data together made the purified cells pool for generating *in*-*silico* mixtures. In total, we have 4 epithelial cell lines, 8 fibroblast lines, 66 B-cells, 336 CD4+ T-cells, and 1277 monocytes.

We also downloaded processed WGBS bed files from IHEC (http://epigenomesportal.ca/ihec) of samples that we expected to be of relatively high purity, representing epithelial, fibroblast and immune cell-subtypes. Samples of low WGBS coverage / sequencing depth were removed. A total of 6 epithelial (4 podocytes and 2 breast epithelial), 2 fibroblast/stromal, 8 B-cells, 11 CD4+ T-cells, and 9 monocyte samples were included. After merging them together, the number of common CpGs was 220351. Only 381 out of the 716 cell-type CpGs in EpiFibIC reference could be found. Thus, purity estimates were checked using 381 CpGs in the EpiFibIC reference using EpiDISH. One podocyte sample was predicted to have a high fibroblast fraction and was removed. For the in-silico mixtures, we thus used 5 epithelial, 2 fibroblast, 8 B-cells, 11 CD4+ T-cells, and 9 monocyte WGBS samples.

### Generation of *in*-*silico* mixtures

To test performance of CellDMC in terms of sensitivity and specificity, we generated 200 *in*-*silico* mixtures with half of them representing controls and the other half representing cases (disease). For cases, we altered the DNAm levels of 150 CpGs in specific cell-types, thus defining true DMCTs (and DMCs).

For each of the 200 mixtures, we randomly sampled an epithelial cell, fibroblast, B-cell, CD4+ T-cell, and monocyte from corresponding pools of purified cells, and mixed them together with 5 weights drawn from a Dirichlet distribution. Since for each sample, DNAm changes were made to all immune cells (B-cells, CD4+ T-cells, and monocytes) simultaneously and equally, we could treat this as a three cell-type mixture (Epi, Fib, and IC) problem, where we inferred the total IC fraction alongside the total epithelial and fibroblast fractions. We considered 5 different DMCT scenarios, including unidirectional DNAm changes in all 3 cell-types, unidirectional changes in 2 cell-types, change in 1 cell-type, bidirectional changes in 3 cell-types, and bidirectional changes in 2 cell-types. For each scenario, we also had 5 different signal to noise ratios (SNR) with values of 3, 2.4, 1.8, 1.2, and 0.9, corresponding to approximate absolute DNAm differences in individual cell-types of 0.42, 0.32, 0.22, 0.12, and 0.12, respectively. The latter two SNR values differ because we increased the noise/variance in the case where SNR=0.9. For every scenario, we ran 100 Monte-Carlo runs. The specific DMCT scenarios for the three main cell-types are summarized in **SI Table.S1**.

We note that we also considered scenarios, where a DNAm change was only induced in a particular IC cell-subtype (and not in all ICs simultaneously), in which case we used HEpiDISH to estimate the cell-type fractions for all IC cell subtypes, in addition to the epithelial and fibroblast fractions. We also considered a scenario mimicking a blood EWAS, where we only mixed together purified IC subtypes.

The overall strategy used to generate in-silico mixtures with a continuous phenotype is similar to that used for a binary phenotype. For each sample, the continuous phenotype y was assumed to take a value between 0 and 1. In the case where y=1, the corresponding beta value was sampled as described previously for disease samples in the binary phenotype context, whereas for samples with values y <1, the altered beta-values were correspondingly scaled by y. The SNR values were defined at y=1.

In the scenario where we allow for a bi-modal distribution within the disease phenotype, we use the same parameters as for the binary phenotype case. The only difference is that for each DMCT, only 70% of the disease samples were randomly selected to be altered, whereas the other 30% disease samples were modelled as control/normal samples.

### Perturbations of cell-type fraction point estimates

We used two different approaches to assess the robustness of CellDMC to the use of cell-type fraction point estimates. Briefly, in one approach, instead of using the point estimates, we randomly sampled a number from the interval point estimate ± standard deviations^∗^3. In the second approach, we generated a bootstrap sample of the DMCs making up the reference matrix, so that a new reference DNAm matrix with the same number of cell-type specific DMC, but with several of these appearing multiple times. Inference of cell-type fractions was then performed using this bootstrapped DNAm reference, with the new point estimates subsequently used in CellDMC.

### Simulations involving missing cell-types in DNAm reference matrix

We considered the case where there are 3 cell-types with one cell-type being differentially methylated (Uni-1C scenario). Specifically, we only altered epithelial cells introducing 150 true epithelial DMCTs. Cell-type fractions were then estimated using the EpiFibIC reference matrix but with one-celltype (fibroblasts) missing from the reference DNAm matrix. Correspondingly, CellDMC was subsequently run with only 2 cell-types. In one case, when generating the *in*-*silico* mixtures, the cell-type fractions for all 3 cell-types were drawn from a uniform Dirichlet distribution, so that each cell-type exhibited the same underlying mean and variance for the fraction.

### Implementation of CellDMC on *in*-*silico* mixtures and definition of sensitivity and specificity

We estimated cell-type fractions of all in-silico mixtures using EpiDISH^8^ (RPC mode) with a reference^17^, which consists of 716 cell-type specific DMCs, to estimate fractions of 3 major cell-types(epithelial cells, fibroblasts, and total immune cells). Then, we run CellDMC with the estimated fractions, multiple hypothesis correction method set to “FDR”, and adjusted P-value threshold 0.05, which resulted in a matrix containing predicted DMCT(s) and a matrix of coefficients for each cell-type including predicted DNAm change, raw and adjusted P-value, ranked by selected cell-type. As long as one cell-type was predicted as DMCT, it was counted as a predicted DMC. Sensitivity of DMC was defined as number of true predicted DMCs divided by 150 (true number of DMCs). Sensitivity of DMCT was defined as the mean sensitivity to correctly predict DMCTs (considering directionality of change) in all cell-types. In the case where we induced DNAm changes in individual IC cell subtypes, we estimated cell-type fractions for all cell-types using our HEpiDISH framework, but all subsequent analysis proceeded as for the 3 cell-type scenario above.

### Power calculations

To estimate the appropriate sample size to achieve a certain sensitivity and specificity, we performed corresponding power calculations for a DNAm change in one cell-type and unidirectional changes in all cell-types scenarios. For each scenario, we did 100 Monte-Carlo runs for 5 SNR(s) as described earlier. We varied sample size from 10 to 500, including 10, 20, 30, 40, 50, 100, 150, 200, 300, and 500, with half of them as controls and half as cases. Mean sensitivities for DMCT detection over 100 Monte-Carlo runs were calculated.

### Cell-type complexity

For K=3, we mixed epithelial, fibroblast and monocytes and used EpiDISH with centEpiFibIC DNAm reference to estimate cell-type fractions. For larger K values, we iteratively added one more IC cell subtype to the mix in the order of neutrophils (K=4), CD4+ T-cells (K=5), B-cells (K=6) and CD8+ T-cells (K=7). For K>3, we used HEpiDISH. For all K, in each Monte-Carlo, there were 150 true DMCTs (75 hyper, and 75 hypo) occurring specifically in the monocytes.

### Application of CellDMC to a blood rheumatoid arthritis EWAS to detect B-cell specific rheumatoid arthritis DMCs

For Rheumatoid Arthritis (RA) EWAS, we obtained Illumina 450k data profiling peripheral blood for 335 controls and 354 cases from Liu et al ^5^ (GSE42861). Data was normalized as described elsewhere ^29^ We estimated cell-type fractions of all 7 major immune cell subtypes (B-cells, CD4T-cells, CD8T-cells, NK cells, monocytes, neutrophils, eosinophils) using EpiDISH with a DNAm reference consisting of 333 IC cell-type specific DMCs ^16^ All parameters of CellDMC were the default choices. We tested CellDMC’s predictions against 10 B-cell specific RA CpGs, which had been validated in 3 independent purified B-cell EWAS datasets, as reported in Julià et al^24^. To assess the statistical significance of CellDMC’s DMCT predictions, we randomly picked 10 CpGs 100,000 times, and compared the weighted t-statistics of the estimated B-cell specific regression coefficients to the observed one. P value was calculated as the number of runs, where the statistic was larger than the original observed one, divided by the number of runs.

### Application of CellDMC to endometrial cancer

We downloaded and processed level 3 Illumina 450k data of uterine corpus endometrial carcinoma(UCEC) from TCGA^12^. This dataset contains 374 cancer samples and 34 normal adjacent control samples. We estimated cell-type fractions for the epithelial, fibroblast and total immune cell components, using our previously validated EpiFibIC DNAm reference (716 cell-type specific DMCs) ^17^.

### Application of CellDMC to a breast cancer EWAS

To test CellDMC in its ability to detect breast cancer epithelial DMCs, we used a breast cancer tissue DNAm 450k dataset, which included 92 normal controls and 305 breast cancer cases, from Teschendorff et al^18^ (GSE69914). We estimated cell-type fractions using EpiDISH in conjunction with a previously validated DNAm reference for breast tissue, consisting of 491 cell-type specific DMCs and 4 cell-types (epithelial, fibroblast, fat and total IC) ^17^. All parameters choices in CellDMC were the default ones. The true positive breast cancer epithelial DMCs and a corresponding list of true negatives were constructed as reported in Zheng et al ^9,17^

### Application of CellDMC to a buccal smoking-EWAS and validation in lung squamous cell carcinoma

The smoking buccal Illumina 450k dataset was processed as described in Teschendorff et al ^30^. Using EpiDISH and EpiFibIC DNAm reference ^17^, we estimated cell-type fractions of epithelial cells, fibroblasts, and total immune cells. Since the estimated fractions of fibroblasts in all samples were quite small with mean less than 0.05, we only included epithelial cells and immune cells when running CellDMC with phenotype of interest as the smoking pack per year. Among all 790 samples in the dataset, we only used 647 samples, of which we had smoking pack per year information available.

DMCTs predicted to occur specifically in the epithelial compartment of buccal swabs were validated in the Illumina 450k lung squamous cell carcinoma (LSCC) data, generated as part of The Cancer Genome Atlas (TCGA) ^32^. We only validated DMCTs which exhibited similar DNAm levels in epithelial and blood cell-subtypes in samples not exposed to smoking, since only for these we would expect to see an association between their DNAm levels and the estimated epithelial fraction in the tumors. To this end, we collected Illumina 450k data of 11 different epithelial cell-lines (Hipe, Saec, Hre, Hae, Hrpe, Prec, Hee, Hcpe, Hnpce, Hmec, Hrce) from ENCODE (GSE40699) and a total of 42 purified samples representing all 7 major immune cell types (neutrophils, eosinophils, monocytes, CD4+ and CD8+ T-cells, B-cells and NK-cells) from Reinius et al ^41^ (GSE35069) to identify CpGs with the same ground state of DNAm level in epithelial and immune cells: in detail, we used limma (an empirical Bayes framework) ^42^, to select 277,801 CpGs with P-values greater than 0.5 or absolute DNAm change between epithelial and immune cells less than 0.05. Of 17,285 and 6658 DMCTs predicted by CellDMC to be hypermethylated and hypomethylated in only epithelial cells, respectively, 3629 and 816 of them overlapped with the 277,801 non-DMCs. We used EpiDISH with EpiFibIC reference to estimate cell-type fractions in the TCGA LSCC DNAm dataset (275 cancer samples). We computed the Pearson’s correlation coefficient between the DNAm beta values of these predicted epithelial DMCTs and the estimated fraction of epithelial cells across the 275 cancer samples. To test whether there is an association between DNAm variance and epithelial cell-type fraction, we used the Breusch-Pagan (BP) test with beta values as response and cell-type fraction as predictor. We used the signed BP-statistic to test for positive and negative associations between DNAm variance with epithelial and immune cell-type fractions, respectively.

### Data availability

All data analyzed in this manuscript is already publicly available from Gene Expression Omnibus - GEO (www.ncbi.nlm.nih.gov/geo/), the TCGA data portal (https://gdc.cancer.gov), International Human Epigenome Consortium - IHEC (http://epigenomesportal.ca/ihec), or from ArrayExpress (https://www.ebi.ac.uk/arrayexpress/). Accession codes for data from GEO include GSE31848, GSE59250, GSE71955, GSE71244, GSE43976, GSE50222, GSE56047, GSE42861, GSE69914, GSE40699, and GSE35069. Accession code for data from ArrayExpress is E-MTAB-2145. All DNA methylation reference matrices used to estimate cell-type fractions are available on Github (https://github.com/sjczheng/EpiDISH). Detailed dataset descriptions and how these data were integrated can be found in Online Methods.

